# Cannabinoid Receptor 2 (CB_2_) Signals via G-alpha-s and Induces IL-6 and IL-10 Cytokine Secretion in Human Primary Leukocytes

**DOI:** 10.1101/663831

**Authors:** Yurii Saroz, Dan T. Kho, Michelle Glass, Euan Scott Graham, Natasha Lillia Grimsey

## Abstract

Cannabinoid receptor 2 (CB_2_) is a promising therapeutic target for immunological modulation. There is, however, a deficit of knowledge regarding CB_2_ signaling and function in human primary immunocompetent cells. We applied an experimental paradigm which closely models the *in situ* state of human primary leukocytes (PBMC; peripheral blood mononuclear cells) to characterize activation of a number of signaling pathways in response to a CB_2_-selective ligand (HU308). We observed a “lag” phase of unchanged cAMP concentration prior to development of classically-expected Gα_i_-mediated inhibition of cAMP synthesis. Application of G protein inhibitors revealed that this apparent lag was a result of counteraction of Gα_i_ effects by concurrent Gα_s_ activation. Monitoring downstream signaling events, activation of p38 was mediated by Gα_i_ whereas ERK1/2 and Akt phosphorylation were mediated by Gα_i_-coupled βγ. Activation of CREB integrated multiple components; Gα_s_ and βγ mediated ∼85% of the response, while ∼15% was attributed to Gα_i_. Responses to HU308 had an important functional outcome – secretion of interleukins 6 (IL-6) and 10 (IL-10). IL-2, IL-4, IL-12, IL-13, IL-17A, MIP-1α, and TNF-α were unaffected. IL-6/IL-10 induction had a similar G protein coupling profile to CREB activation. All response potencies were consistent with that expected for HU308 acting via CB_2_. Additionally, signaling and functional effects were completely blocked by a CB_2_-selective inverse agonist, giving additional evidence for CB_2_ involvement. This work expands the current paradigm regarding cannabinoid immunomodulation and reinforces the potential utility of CB_2_ ligands as immunomodulatory therapeutics.

**Significance statement:** Cannabinoid receptor 2 (CB2) is a G protein-coupled receptor which plays a complex role in immunomodulation and is a promising target in a range of disorders with immune system involvement. However, to date the majority of the studies in this field have been performed on cell lines, rodent models, or stimulated primary cells. Here we provide a detailed account of CB2-mediated signaling in primary human immune cells under conditions which closely mimic their *in vivo* state. We reveal a complex signaling system involving an unprecedented CB2 signaling pathway and leading to immunomodulatory functional outcomes. This work provides not only a critical foundation impacting CB2-targeted drug discovery, but reveals important wider considerations for GPCR signaling studies and model validity.

**Table of Contents Summary Figure:** 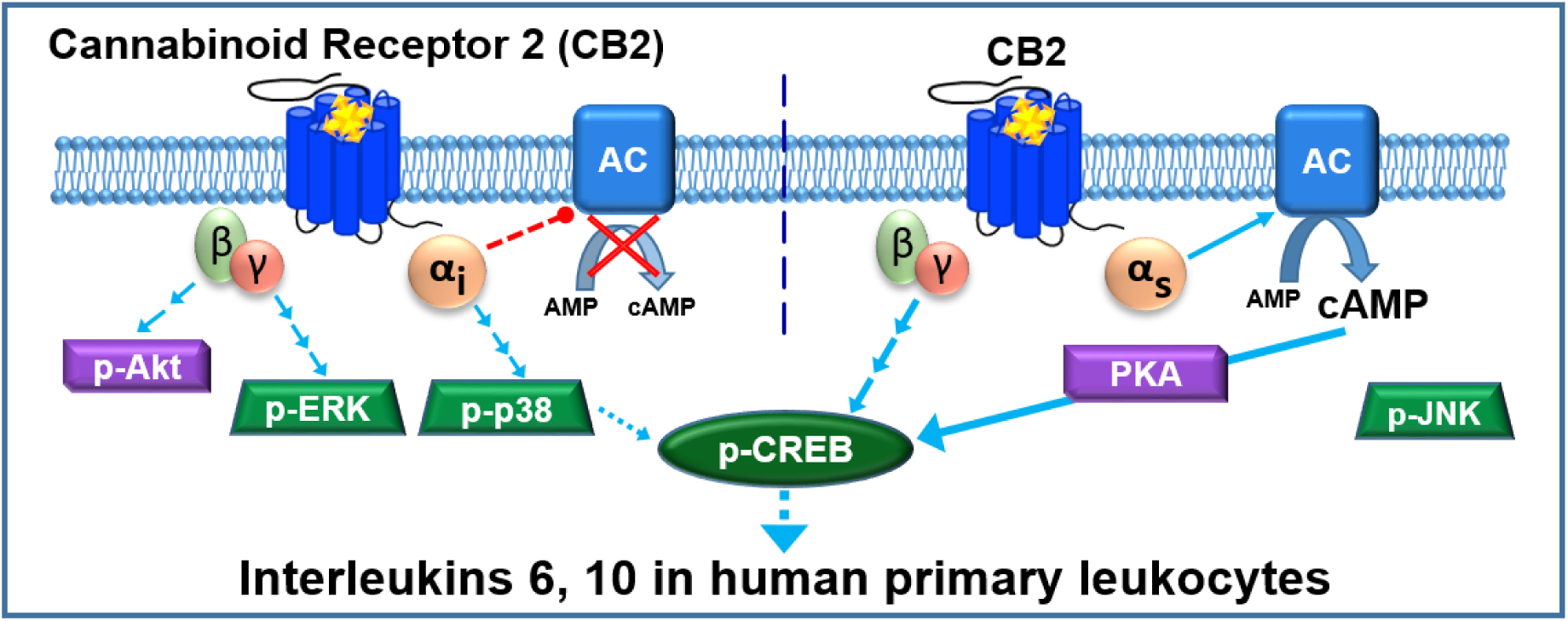

## Introduction

Cannabinoid Receptor 2 (CB_2_) is a class A G protein-coupled receptor (GPCR) which is expressed primarily in the immune system ^e.g. 1,2^ and is a promising therapeutic target for immune modulation in a wide range of disorders ^reviewed in 3^. CB_2_ couples to Gα_i/o_ proteins inhibiting adenylyl cyclases ^e.g. 4,5^, and limited evidence indicates coupling to Gα_q_ ^e.g. 6^. To date, there is no evidence for CB_2_ coupling to Gα_s_; although increased cyclic adenosine monophosphate (cAMP) synthesis has been reported, this was coupled to Gα_i/o_, likely via Gβγ ^7,8^. Downstream of G protein coupling, and perhaps involving modulation by β-arrestins ^9^, activation of extracellular signal-regulated kinases 1 and 2 (ERK1/2) is a widely studied consequence of CB_2_ activation reported in cell lines heterologously expressing this receptor ^e.g. 10^ and in rodent ^e.g. 11^ and human immunocompetent cell lines ^e.g. 8,12^, with limited published evidence for such signaling in human primary leukocytes ^13,14^. The paucity of studies on human primary leukocytes may in part be due to methodological difficulties such as low cell yields. Mitogenic or antigenic proliferative stimulations are widely used approaches to overcome some of these practical limitations ^e.g. 13,14^. However, these methods cause lymphocytes to de-differentiate and turn into lymphoblasts – mitotically overactive cells – which do not closely model the *in situ* state of leukocytes ^15^.

The limited literature regarding CB_2_ signaling via other types of mitogen-activated protein kinases (MAPKs) indicates context-dependence of activation. CB_2_ can induce p38 phosphorylation (p-p38)^e.g. 16^, which has been linked to growth inhibition of cancer cells, yet in LPS-stimulated monocytes p-p38 was decreased by a CB_2_-selective agonist ^12^, while B lymphocytes ^17^ and dendritic cells ^18^ were non-responsive. Akt kinase (protein kinase B), an important mediator of immunomodulation ^reviewed in 19^ can be activated ^20^, inhibited ^21^, or unaffected ^22^ by CB_2_ activation. A stress-activated protein kinase JNK (c-Jun NH2-terminal kinase), involved in pro-inflammatory signaling ^reviewed in 23^, has been shown to be activated ^24,25^ and inhibited ^12^ via CB_2_.

The ultimate outcomes of CB_2_ activation in immune cells include inhibition of proliferation, induction of apoptosis, influences on cytokine/chemokine networks, and regulation of adhesion and migration ^reviewed in 26^. In human immunocompetent cells CB_2_ activation inhibits IL-2 ^e.g. 27^, IL-17, INF-γ, TNF-α ^e.g. 28^ secretion, and stimulates IL-4 ^29^ and TGF-β ^30^ secretion. Murine *in vitro* and *in vivo* models indicate that cannabinoids inhibit IL-2 ^31^, IL-12 and IFN-γ ^32,33^, and induce IL-4 secretion ^32,33^. However, these studies stimulated leukocytes with mitogens or antibodies. Species differences in cannabinoid signaling notwithstanding ^e.g. 5^, cytokine secretion is generally highly dependent on cellular environment. Indeed, cannabinoid effects on the cytokine network are sensitive to the method of cell activation ^34^.

Therefore, close to the entirety of our current understanding of CB_2_ signaling and influence on the cytokine secretome has been obtained from models which exhibit considerable limitations in modelling normal human physiology. We endeavored to carry out a comprehensive study of CB_2_ signaling in unstimulated human primary leukocytes (peripheral blood mononuclear cells, PBMC) under conditions closely preserving their *in vivo* state. We have observed an unprecedented CB_2_-mediated signaling and functional profile, including coupling to Gα_s_.

## Results

### CB_2_, but not CB_1_, is expressed in unstimulated human primary PBMC

We quantified expression of CB_1_ and CB_2_ protein by whole cell radioligand binding with high affinity CB_1_/CB_2_ ligand [^3^H]-CP55940 displaced by CB_1_- and CB_2_-selective ligands, ACEA ^35^ and HU308 ^36^ respectively. Although CB_2_ was expected to be the primary cannabinoid-responsive receptor in human PBMC, CB_1_ transcripts have also been detected in some contexts ^e.g. 1^. Whereas CB_1_ protein could not be detected in PBMC from any of three healthy human donors, CB_2_ was readily detectable (Fig. 1). Intra-subject variability in CB_2_ expression was low (independent experiments are from three blood draws from the same donor taken over the course of up to 8 months), while inter-subject expression was within the same magnitude, ranging from 704 to 1323 fmol/mg. PBMC samples from these same three donors were utilized in subsequent experiments.

**Fig. 1.**
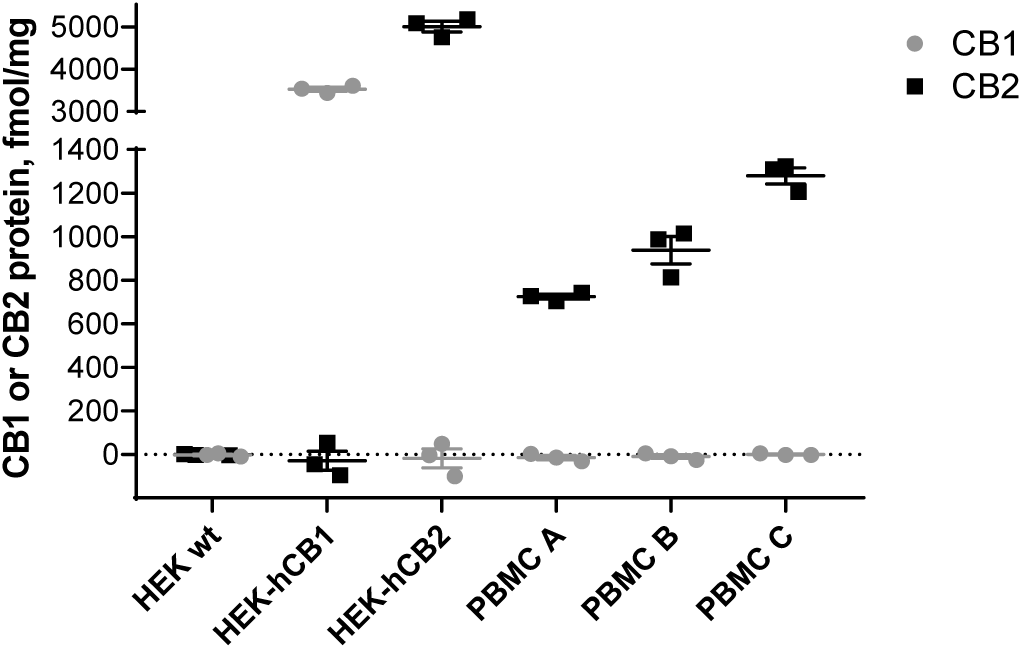
CB_1_ and CB_2_ protein quantification by whole cell radioligand binding. Cells were incubated with [^3^H]-CP55940 (5nM) in the presence and absence of a CB_1_-selective displacer ACEA (1µM), or a CB_2_-selective displacer HU308 (1µM). Total binding sites (B_max_) were calculated (see methods) and converted to fmol/mg of total protein. HEK wt is a negative control (not expressing CB_1_ or CB_2_), HEK-hCB_1_ is a positive control for CB_1_, HEK-hCB_2_ is a positive control for CB_2_. PBMC A, B, and C are from subjects A, B, and C, respectively. The graph shows independent experiment means (from technical triplicate) as well as overall mean ± SEM of these three independent experiments each performed with cells from a separate subject (three subjects in total).

### CB_2_ activation induces simultaneous Gα_i_ and Gα_s_ coupling, producing delayed cAMP flux

CB_2_ is widely recognized to inhibit cAMP production via Gα_i/o_ inhibition of adenylyl cyclases. Depending on basal adenylyl cyclase activity in the sample of interest, stimulation of adenylyl cyclase (e.g. with forskolin) is a typical prerequisite for measuring inhibition ^4,10^. We first studied the time-course of cAMP signaling in PBMC in response to CB_2_-selective agonist HU308 (1µM), and observed inhibition of forskolin-stimulated cAMP synthesis in line with our general expectations (Fig. 2A). However, there was a lag to the onset of measurable inhibition which lasted at least 10 min, with the 20-35 min time-points being significantly different from time-matched vehicle control (*p < 0.01*). This was followed by a steady reduction in cAMP concentration, reaching a maximum extent by 30 min (to 60.3 ± 2.8% of forskolin-stimulated), with cAMP subsequently returning back to the forskolin-induced level by 60 min. Prior studies in cells overexpressing CB_2_ describe an immediate onset and earlier peak cAMP inhibition ^37^. Given that we cannot readily measure inhibition of cAMP production without forskolin being present, we hypothesized that low basal cAMP and/or slow response to forskolin in the primary leukocytes might mask the ability to detect inhibition at early time-points and hence produce the observed delay. However, pre-incubation with forskolin prior to HU308 addition did not influence the signaling latency, time to peak response, nor efficacy, therefore ruling out this hypothesis (Fig. S1).

**Fig. 2.**
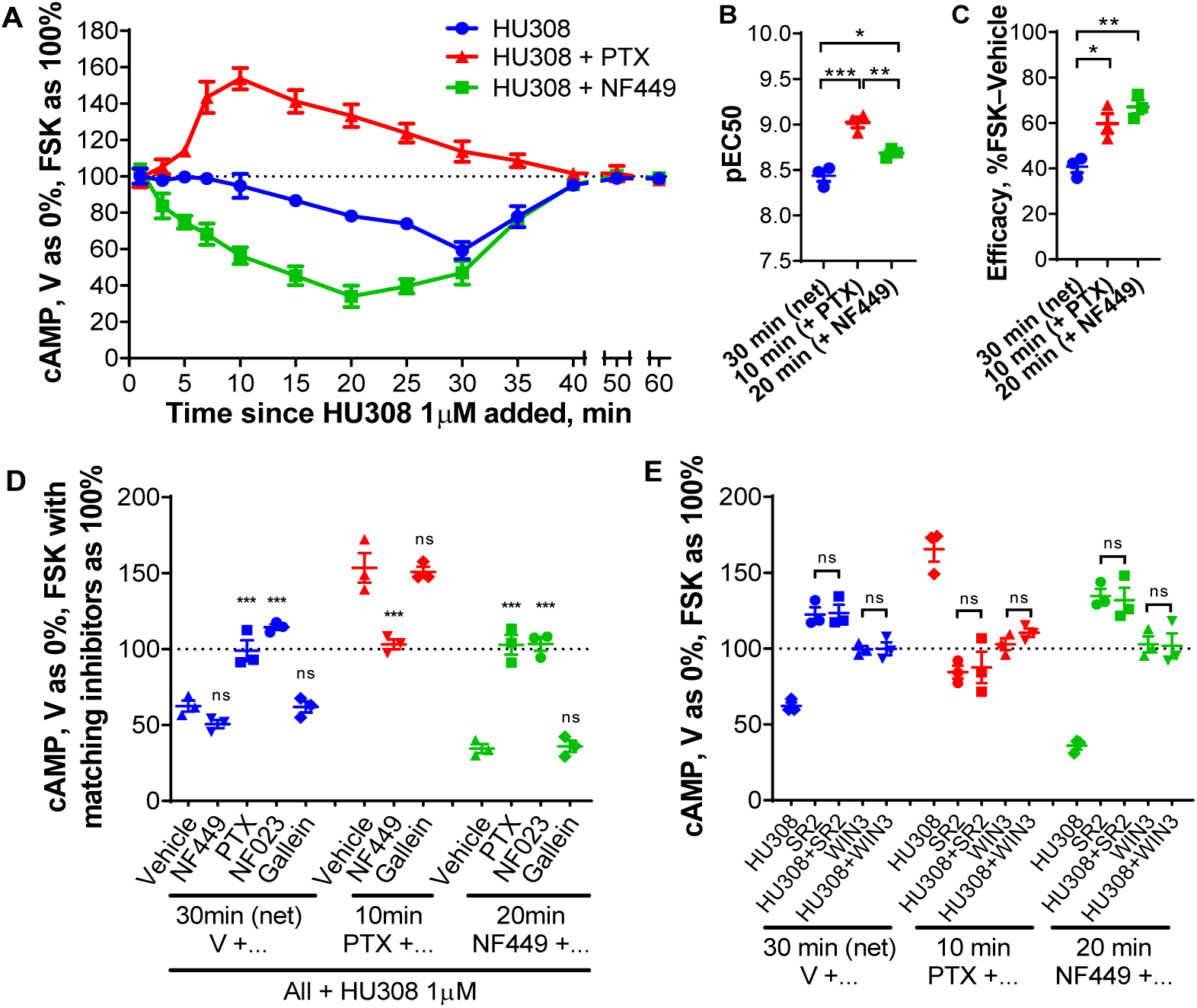
cAMP signaling in human primary PBMC. **A.** PBMC were incubated with 10µM forskolin (FSK) and HU308 (1µM) without inhibitors (○), or after pre-treatment with NF449 (10µM for 30 min, ▪), or with PTX (100ng/mL for 6 h, ▴). The graph shows mean ± SEM of three independent experiments performed in technical triplicate, each with cells from a separate subject (three subjects in total). **B, C.** HU308 concentration-response curve parameters; potency (**B**) and efficacy (**C**; span of curves as in Fig. S3)**. D, E**. Stimulations were with 10µM forskolin and 1µM HU308 or vehicle for 30 min for net cAMP flux, 10 min for the stimulatory pathway (revealed by PTX), or 20 min for the inhibitory pathway (revealed by NF449). **D.** PBMC were additionally pre-treated as indicated with PTX, NF449, NF023 or gallein. Data are normalized to vehicle control as 0% and forskolin controls with matching inhibitors as 100%. Statistical comparisons are to the vehicle plus HU308 control within each set. **E.** PBMC were pre-treated as indicated, then co-incubated with forskolin (10µM), SR144528 (SR2, 1µM) or WIN55212-3 (WIN-3, 5µM), and HU308 (100nM) or vehicle for 10, 20, or 30 min. Data are normalized to vehicle control as 0% and forskolin control as 100%. All SR2 and WIN-3 conditions were significantly different from “HU308” control for peak signaling within each set (*P < 0.001*). Within each set, condition “SR2” was compared to “SR2 + HU308”, “WIN3” was compared to “WIN3 + HU308”, and had no significant difference. Graphs **B-E** show independent experiment means (from technical triplicate) as well as overall mean ± SEM of these three independent experiments each performed with cells from a separate subject (three subjects in total).

Pre-incubation with pertussis toxin (PTX), a Gα_i_ ^e.g. 38^ and Gα_i_-derived βγ ^reviewed in 39^ inhibitor, followed by its co-application with HU308 revealed a wave of increased cAMP (Fig. 2A). This induction of cAMP accumulation had an earlier onset than the net cAMP inhibition (without PTX), with peak activation at around 10 min, and a slower return to forskolin-stimulated levels than the net cAMP flux, with the 7-25 min time-points being significantly different from time-matched vehicle control (*p < 0.01*). Given that increased cAMP is associated with Gα_s_ coupling, we tested the effect of Gα_s_-blocker NF449 ^e.g. 40^ on HU308-stimulated cAMP flux. In comparison with net cAMP flux, not only was the peak of inhibition shifted to an earlier time-point (20 min), but the onset of cAMP inhibition had no apparent delay, with the 3-35 min time-points being significantly different from time-matched vehicle control (*p < 0.01*). The time-courses for onset of reduction and return to forskolin-stimulated cAMP levels for this now apparently “pure” inhibitory signaling had similar timing.

These findings indicate that when no signaling inhibitors are present the opposing influences of Gα_i_ and Gα_s_ coupling produce a net apparent “lag” to the onset cAMP flux. At later time-points, the different time-courses and relative efficacies of Gα_i_ versus Gα_s_ coupling to adenylyl cyclase(s) result in transient net inhibition of cAMP synthesis. Increased cAMP in response to HU308 was detectable even without forskolin stimulation, although inhibition of Gα_i_ was required to reveal this as was the case when forskolin was present (Fig. S2).

cAMP fluxes for the three types of responses at their respective peak time-points (net cAMP, 30 min; cAMP accumulation, 10 min; cAMP inhibition, 20 min) were of nanomolar potency, typical for CB_2_-mediated signaling by HU308 ^5,41^, with subtle differences in potencies between the three cAMP responses with rank order: stimulatory cAMP > “pure” inhibitory > net cAMP (Fig. 2B, S3). Not surprisingly from the opposing effects on cAMP concentration, the peak stimulatory and “pure” inhibitory cAMP responses revealed by the Gα inhibitors had greater efficacies than the maximal net cAMP response (Fig. 2C, S3). Between-subject variability was small.

We further probed the contribution of G proteins (Fig. 2D) and investigated CB_2_-specificity (Fig. 2E). As expected from the time-course data, Gα_s_ inhibitor NF449 had a minimal effect on the net cAMP response peak (30 min), while Gα_i/o_/βγ inhibitor PTX and Gα_i_ inhibitor NF023 ^e.g. 42^ blocked the HU308-induced inhibition of cAMP synthesis. The “pure” inhibitory response (in the presence of NF449) was also completely blocked by the Gα_i/o_ inhibitors. In contrast, the peak stimulation of cAMP synthesis (measured in the presence of PTX) was completely blocked by Gα_s_ inhibitor NF449. A Gβγ inhibitor gallein ^e.g. 43^ had no effect on cAMP signaling, indicating lack of Gβγ involvement (Fig. 2D). As well as the potencies of HU308-induced cAMP flux being consistent with CB_2_-specific responses, inverse agonist SR144528 ^44^ at 1µM (a concentration at which it binds to CB_2_, but not to CB_1_, Fig. S4) completely blocked cAMP signaling induced by HU308 (Fig. 2E). Inverse agonism of SR144528 was readily measurable for the inhibitory and net cAMP pathways *(p < 0.001)*, in which it increased cAMP above forskolin-induced levels. There was a trend toward inverse agonism in the stimulatory cAMP pathway, but this reduction in cAMP was not significantly different from forskolin alone *(p = 0.78).* A CB_1_/CB_2_ neutral antagonist WIN55212-3 ^45,46^ (no significant differences from forskolin control, *p < 0.05*) also completely abrogated all HU308-induced cAMP signaling (Fig. 2E). Given the lack of CB_1_ protein in PBMC (Fig. 1), WIN55212-3 most likely acted via CB_2_, and taken together with the SR144528 data and response potencies provides strong evidence that the observed signaling is CB_2_-mediated.

### CB_2_-stimulated phosphorylation of ERK1/2 is mediated by Gα_i_-but not Gα_s_-coupled Gβγ

Phosphorylation of ERK1/2 (p-ERK) is a widely observed CB_2_ response, but the potential pathways to stimulation are many and varied and p-ERK can act as an integrator of responses ^reviewed in 47^. Time-courses with 1µM HU308 revealed transient activation of p-ERK, with a peak at around 3 min (Fig. 3A). HU308 activated ERK1/2 in a concentration-dependent manner (Fig. S5), with approximately one log unit lower potency than observed for cAMP fluxes (pEC50 7.69 ± 0.06). Responses were very consistent between the three donors. Neither Gα_i_ inhibitor NF023 nor Gα_s_ inhibitor NF449 had an effect on the observed ERK1/2 signaling (Fig. 3B), whereas Gα_i/o_/βγ inhibitor PTX and Gβγ inhibitor gallein both completely blocked activation of p-ERK. This indicates that p-ERK1/2 induced by HU308 is mediated by Gα_i_-linked βγ dimers. We further verified that the observed p-ERK1/2 response was CB_2_-mediated by co-applying HU308 with SR144528 or WIN55212-3 (Fig. 3C). Both antagonists completely blocked p-ERK, supporting the inference that HU308 binding to CB_2_ receptors is the only initiator of the measured p-ERK signal. Furthermore, we observed no p-ERK response for CB_1_-selective agonist ACEA at 1µM (Fig. S6), a concentration which binds to CB_1_ but not CB_2_ (Fig. 1) and can induce CB_1_-mediated p-ERK ^e.g. 48^.

**Fig. 3.**
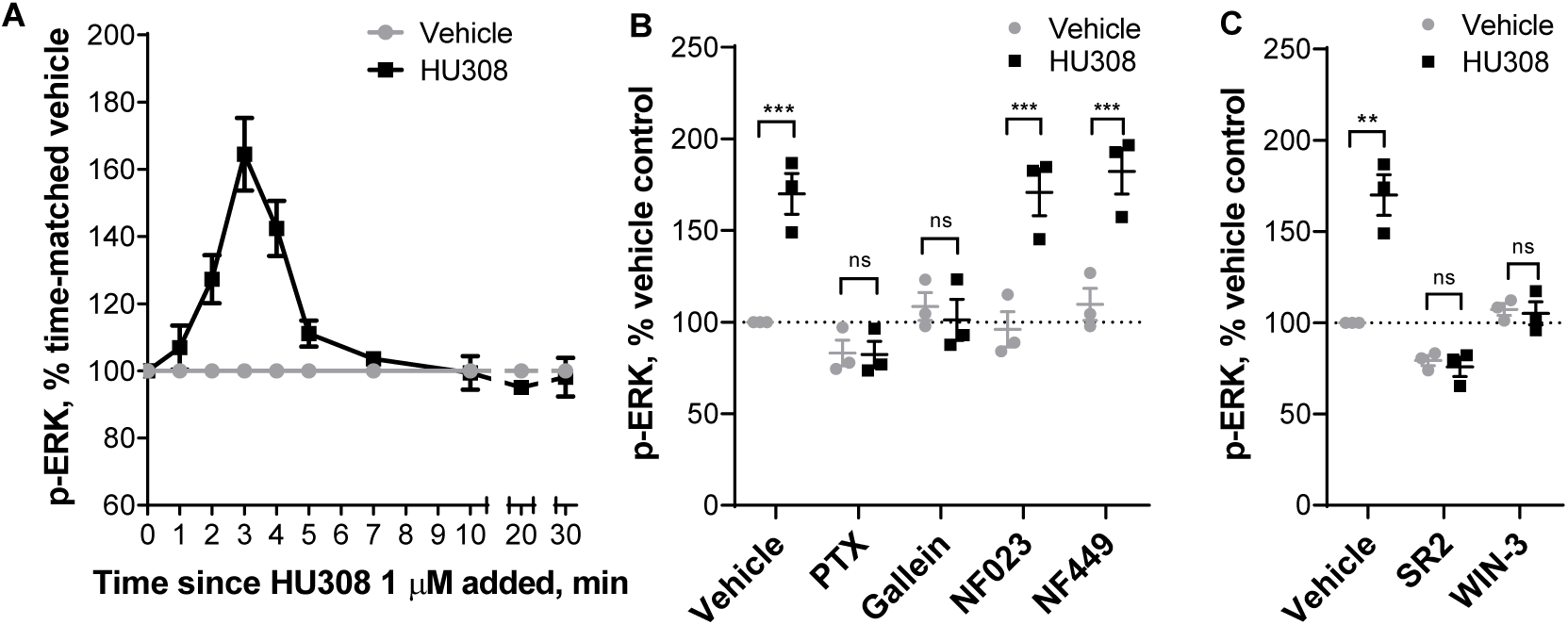
ERK1/2 signaling in human primary PBMC. **A.** Time-course with 1µM HU308 (▪) or a vehicle control (•). 2, 3, and 4 min time-points are significantly different from time matched vehicle controls (*p < 0.01*). The graph shows mean ± SEM of three independent experiments performed in technical triplicate, each with cells from a separate subject (three subjects in total). **B.** 3 min incubation with HU308 (1µM) or vehicle control in the presence of G protein inhibitors PTX, gallein, NF023, NF449 or vehicle control. **C.** 3 min incubation with HU308 (100nM) or vehicle control in the presence of SR144528 (SR2, 1µM), WIN55212-3 (WIN-3, 5µM), or vehicle control. Data are normalized to vehicle control (100%). Graphs **B**,**C** show independent experiment means (from technical triplicate) as well as overall mean ± SEM of these three independent experiments each performed with cells from a separate subject (three subjects in total).

### CB_2_ induces phosphorylation of p38 via Gα_i_ and Akt via Gβγ, but does not activate JNK

Time-courses stimulating PBMC with 1µM HU308 revealed phosphorylation of Akt and p38 with signaling peaks at around 20 min, and with similar overall kinetics (Fig. 4A). Conversely, there was no phosphorylation of JNK detected. Application of PTX completely abrogated Akt and p38 phosphorylation (Fig. 4B), implicating the involvement of Gα_i_/βγ proteins. Gβγ inhibitor gallein completely blocked p-Akt, but did not influence p-p38 (Fig. 4B). HU308 therefore induces phosphorylation of p38 downstream of Gα_i_ subunits, while Akt is activated via Gα_i_ -coupled βγ heterodimers.

**Fig. 4.**
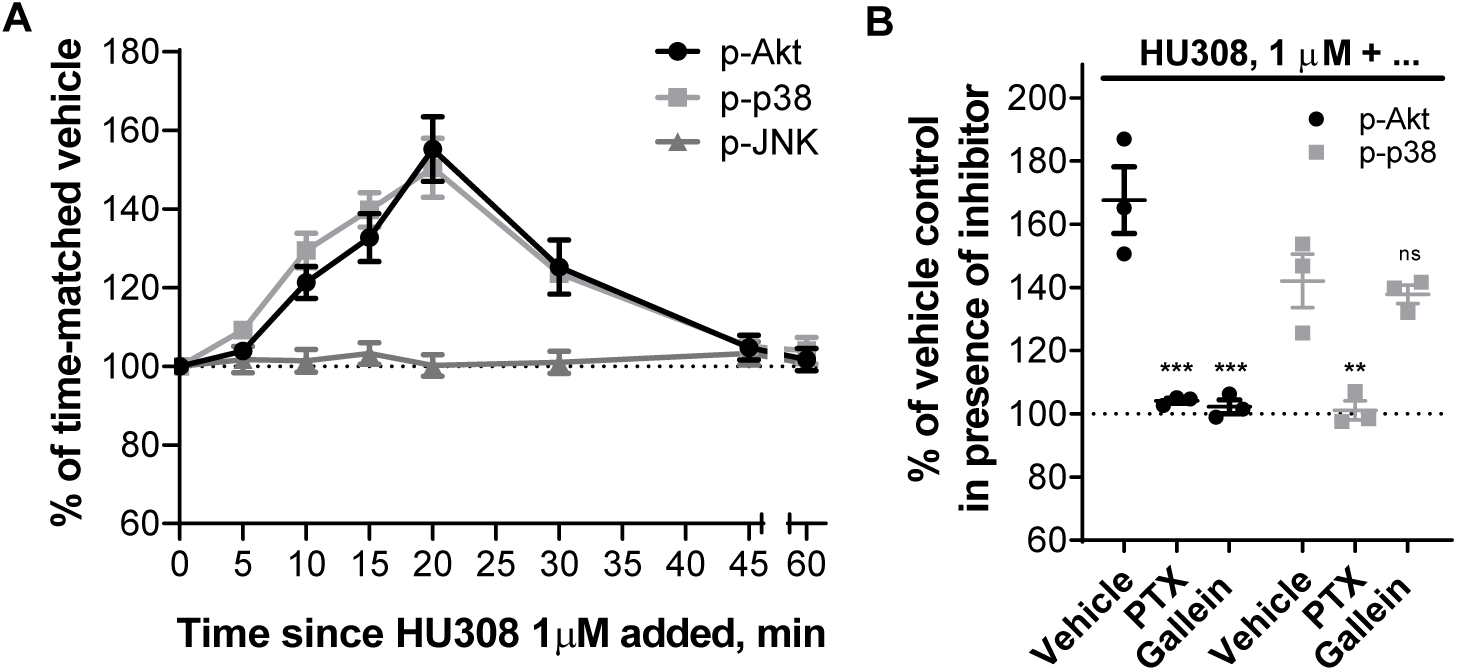
Phosphorylation of Akt, JNK, and p38 MAP kinases in human primary PBMC. **A.** Time-courses with HU308 (1µM) and vehicle control. For p-Akt and p-p38 time-points 10-30 min are significantly different from time-matched vehicle controls (*p < 0.05*). For p-JNK, all points are indistinguishable from 0 min. The graph shows mean ± SEM of three independent experiments performed in technical triplicate, each with cells from a separate subject (three subjects in total). **B.** Responses to 20 min 1µM HU308 after pre-treatment with PTX, gallein or vehicle control at. Data are normalized and statistically compared to the corresponding vehicle plus HU308 control. The graph shows independent experiment means (from technical triplicate) as well as overall mean ± SEM of these three independent experiments each performed with cells from a separate subject (three subjects in total).

### CB_2_ induces p-CREB downstream of Gα_s_ and Gβγ

CREB phosphorylation is a classical downstream consequence of Gα_s_ and cAMP-mediated activation of protein kinase A (PKA) ^reviewed in 49^. A time-course with PBMC stimulated by 1µM HU308 (Fig. 5A) revealed a wave of CREB phosphorylation which was maximal around 30-40 min and resolved by 60 min. We applied G protein inhibitors NF023, NF449, gallein and PTX (Fig. 5B), and found that both NF449 and gallein each partially inhibit CREB phosphorylation to a similar degree, but almost completely block the HU308 response when co-applied indicating that Gα_s_ and βγ heterodimers are similarly important in CREB phosphorylation in this context. p-CREB was reduced by PTX and NF023 to a lesser extent, with equivalent efficacy to each other, implying a minor involvement of Gα_i_ but not Gα_i_-coupled βγ in p-CREB activation. SR144528 and WIN55212-3 completely blocked the peak CREB phosphorylation (Fig. 5C), indicating that this pathway activation occurs downstream of CB_2_. SR144528 reduced p-CREB relative to the vehicle control indicating that this compound acted as an inverse agonist (*p < 0.05*).

**Fig. 5.**
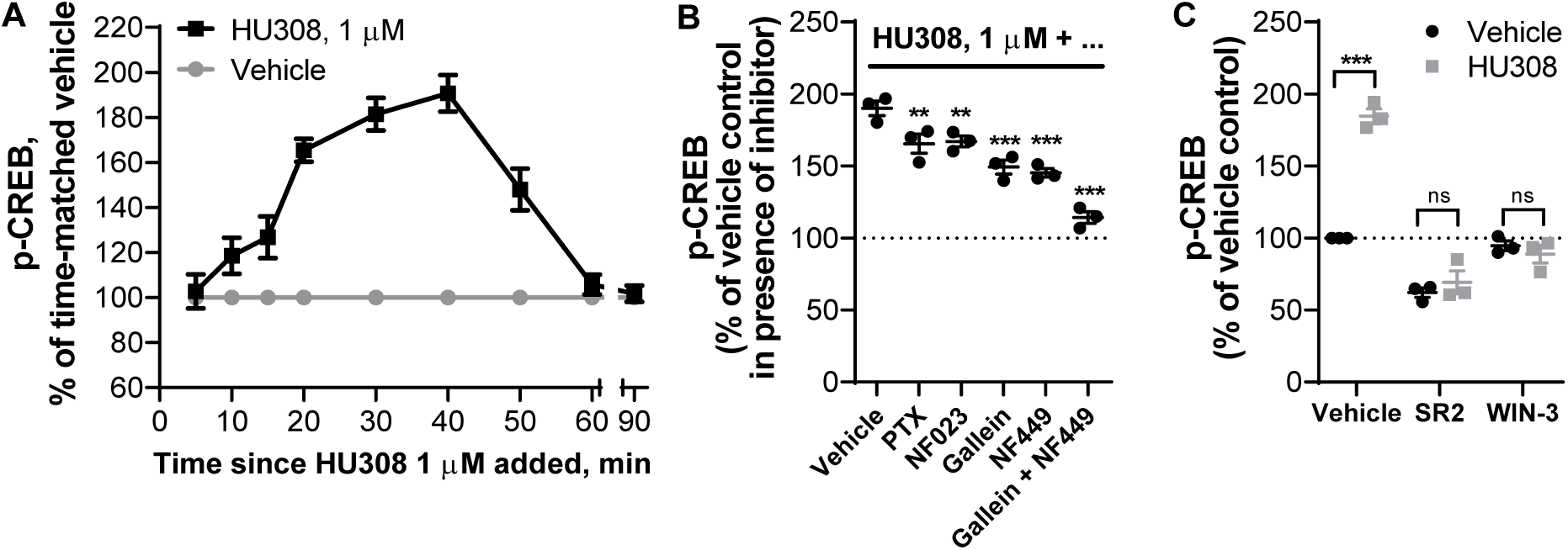
Phosphorylation of CREB in human primary PBMC. **A.** Time-course with HU308 (1µM) and vehicle control. 10-50 min time-points are significantly different from time-matched vehicle controls (*p < 0.01*). The graph shows mean ± SEM of three independent experiments performed in technical triplicate, each with cells from a separate subject (three subjects in total).**B.** p-CREB stimulated by HU308 (1µM) for 40 min in the presence of G protein inhibitors gallein, NF023, NF449, PTX or vehicle control; data are normalized to responses in the absence of HU308 (100%); statistical comparisons are to the HU308 response without inhibitors. **C.** p-CREB stimulated by a CB_2_-selective agonist HU308 (100nM) or vehicle control at 40 min in the presence of a CB_2_-selective inverse agonist/antagonist SR144528 (SR2, 1µM), a CB1/CB2 neutral antagonist WIN55212-3 (WIN-3, 5µM), or vehicle control; data are normalized to vehicle control (100%). Graphs **B**,**C** show independent experiment means (from technical triplicate) as well as overall mean ± SEM of these three independent experiments each performed with cells from a separate subject (three subjects in total).

### CB_2_ activation induces secretion of cytokines IL-6 and IL-10

We stimulated PBMC with HU308 under the same conditions as used for the signaling assays and detected cytokines released into the assay media. After 12 hours, interleukin 6 (IL-6) and 10 (IL-10) concentrations were significantly different from vehicle-treated cells (*p < 0.01*), whereas the remaining 7 cytokines in our panel (IL-2, IL-4, IL-12, IL-13, IL-17A, MIP-1α, and TNF-α) were indistinguishable from vehicle-treated cells (Fig. S7). HU308 was also co-applied with forskolin (10µM) to mimic conditions of our cAMP signaling assays; this had no effect by itself and did not influence the HU308-mediated IL-6 and IL-10 levels (Fig. S7). The responses were very similar between PBMC from 3 subjects and cell numbers were unaffected by the applied ligands (Fig. S8). Subsequent interleukin experiments were carried out *without* forskolin present.

IL-6 and IL-10 accumulation were also detectable at an earlier time-point – after 6 h of HU308 stimulation (Fig. 6A). This was concentration-dependent, with HU308 having greater potency for inducing IL-10 (pEC50 10.06 ± 0.08 and 9.55 ± 0.04; *p = 0.005*, unpaired t test). Application of G protein inhibitors revealed that HU308-induced IL-6 and IL-10 secretion are almost fully blocked by Gα_s_ inhibitor NF449, partially blocked by βγ inhibitor gallein, and to a lesser extent sensitive to Gα_i_ inhibitor NF023 (Fig. 6B, 6C). Responses to HU308 (10nM) were fully abrogated by SR144528 (1µM) and WIN55212-3 (5µM) down to levels below the assays’ limits of quantitation. SR144528 and WIN55212-3 had no measurable effects on IL-6 or IL-10 production when applied alone.

**Fig. 6.**
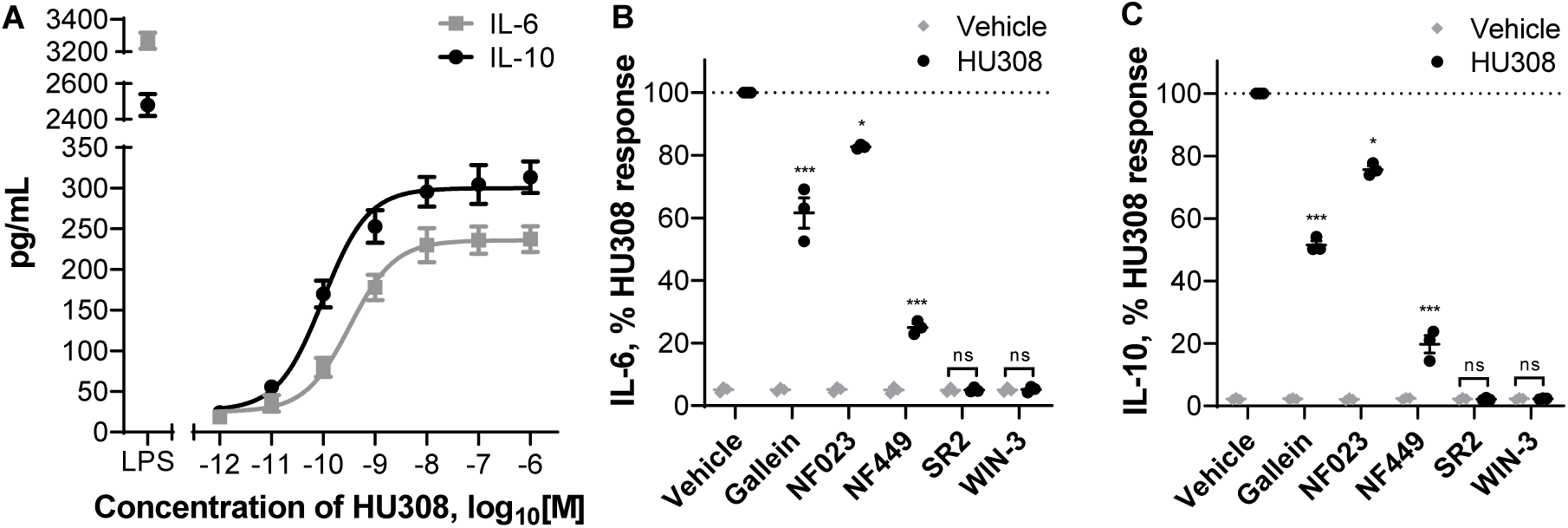
Induction of cytokine secretion by PBMC. **A.** Induction of IL-6 and IL-10 secretion in response to 6 h incubation with HU308. Lipopolysaccharide (LPS, 10ng/mL) is a positive control. The graph shows mean ± SEM of three independent experiments performed in technical triplicate, each with cells from a separate subject (three subjects in total). **B.** IL-6 and IL-10 accumulation in the presence of G protein inhibitors or CB2 antagonists: PBMC were pre-treated with gallein, NF023, NF449, SR144528 (SR2, 1µM), or WIN55212-3 (WIN-3, 5µM) for 30 min followed by 6-hour treatments with HU308 (10nM) with the inhibitors/antagonists present for the entire duration of incubation; data are normalized to HU308 responses without inhibitors/antagonists. Effects of inhibitors on HU308 responses were statistically compared with the corresponding HU308 response without inhibitor. In the presence of SR2 and WIN-3, there was no statistical difference between HU308 and the HU308 vehicle control. Graphs **B**,**C** show independent experiment means (from technical triplicate) as well as overall mean ± SEM of these three independent experiments each performed with cells from a separate subject (three subjects in total).

## Discussion

To our knowledge this study represents the first investigation of CB_2_ signaling and functional implications thereof in unstimulated human primary PBMC. Cells were kept in culture for as little time as practically possible (up to 6 hours prior to stimuli of interest) and in 10% serum in order to mimic the state of leukocytes *in vivo*; a unique experimental paradigm in comparison with the vast majority of other cannabinoid studies to date. Obtaining an accurate understanding of CB_2_ function in normal cells is critical to meaningful interpretation of CB_2_ function in abnormal conditions and in assessing therapeutic potential. The signaling profile we observed is summarized in Fig. 7.

**Fig. 7.**
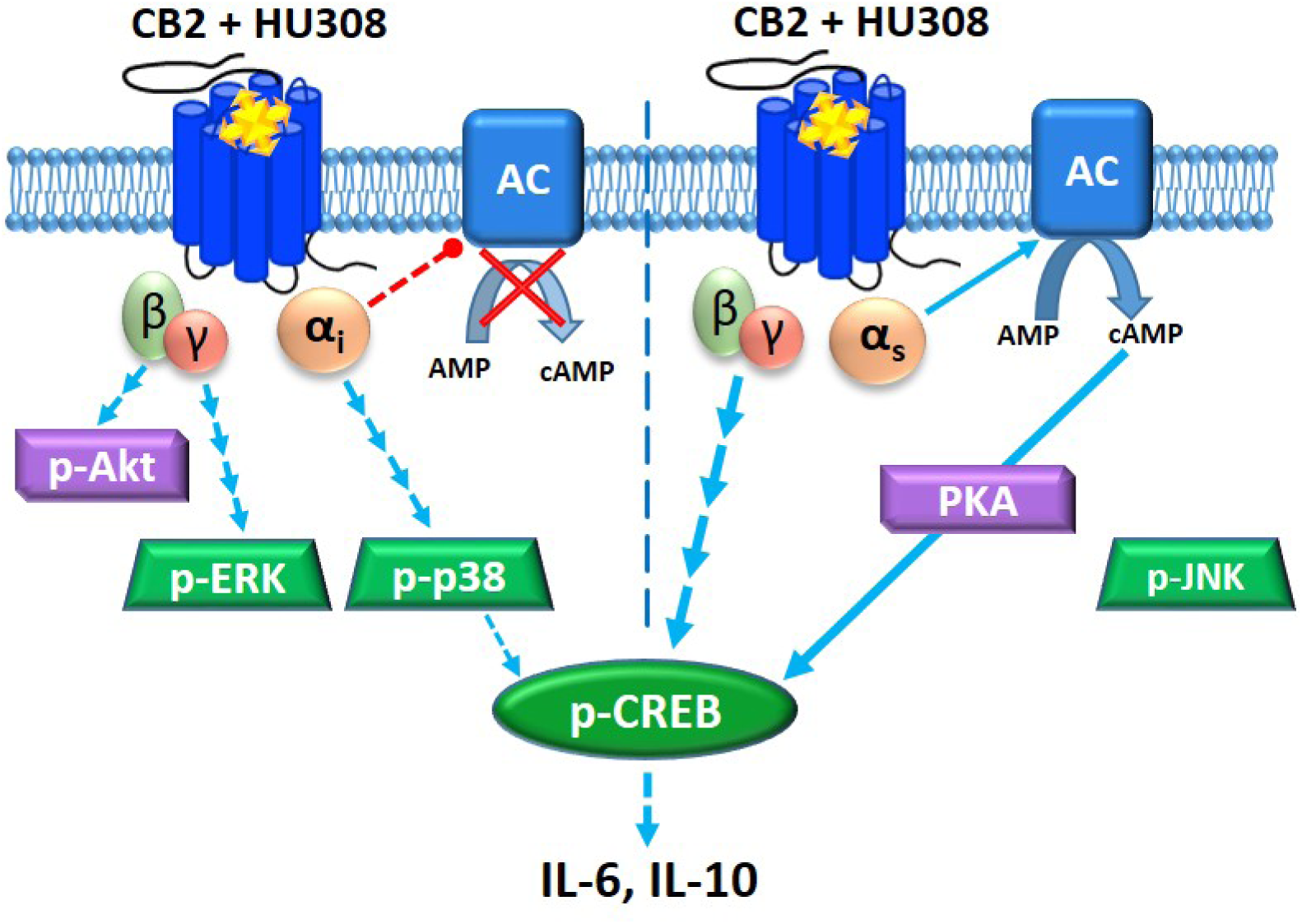
CB_2_ signaling network in mixed human primary leukocytes. Binding of HU308 to CB_2_ leads to activation of Gα_i_βγ and Gα_s_βγ heterotrimers. Gα_i_ inhibits adenylyl cyclase(s) (AC) and activates a cascade leading to phosphorylation of p38 and CREB. βγ from Gα_i_βγ activates Akt and initiates a cascade leading to phosphorylation of ERK. Gα_s_ stimulates AC which leads (likely via protein kinase A, PKA) to p-CREB. βγ from non-Gα_i_βγ (likely from Gα_s_βγ) also contributes to p-CREB activation. Based on inhibitor effects (Fig. 5B), Gα_s_ and non-Gα_i_βγ are the major mediators of p-CREB, whereas p-p38 is a minor p-CREB mediator. JNK phosphorylation was not observed. Given that Gα_i_-mediated inhibition of AC does not inhibit p-CREB activation it is likely that Gα_i_ and Gα_s_ modulation of AC are either segregated in separate compartments within one cell, or occur in different cells (represented by vertical dotted line). p-CREB likely mediates IL-6/IL-10 induction, which might be released from one cell or from different cell types.

We found that human primary PBMC respond in a concentration-dependent manner to CB_2_-selective agonist HU308 with net inhibition of forskolin-induced cAMP synthesis, however a surprising feature was a latency of approximately 10 min prior to any measurable change in cAMP concentration. Application of Gα inhibitors revealed that CB_2_ simultaneously activates stimulatory and inhibitory cAMP pathways, explaining the apparent delay in net cAMP inhibition which arises from these oppositely-directed responses balancing each other at early time-points. Response potencies were consistent with CB_2_ specificity and signaling was also sensitive to SR144528 and WIN55212-3, indicating that the observed signaling is CB_2_-mediated. Although the idea that CB_2_ may couple promiscuously to G proteins is not unprecedented ^6,10^, to our knowledge CB_2_ coupling to Gα_s_ has not been reported previously. Börner and colleagues ^8^ observed stimulatory effects of CB_1_/CB_2_ mixed agonists JWH015, THC and methanandamide on cAMP concentrations in IL-4-stimulated human primary T lymphocytes after prolonged cannabinoid incubations, however cAMP increases were sensitive to PTX and therefore concluded to be downstream of Gα_i_, likely via βγ ^8^. Promiscuous coupling of other receptors to Gα_i_ and Gα_s_ has been observed, including CB_1_ in our own hands ^50^. However, we did not observe Gα_s_ coupling of human CB_2_ in this same model. Studies in other models have also applied PTX and not revealed CB_2_ signaling consistent with Gα_s_ coupling ^e.g. 10,51,52^. We therefore speculate that a critical component of the CB_2_ immune cell signaling landscape has not been captured in previously studied models.

As we have studied mixed PBMC we cannot provide direct evidence as to whether both cAMP inhibition and stimulation have been induced in the same cell or if our findings arise as a result of assaying a mixture of Gα_i_-predominant versus Gα_s_-predominant cell types. Leukocytes are known to express Gα_i_ ^e.g. 53,54^, Gα_s_ ^reviewed in 39^, and Gα_i/s_-sensitive adenylyl cyclases ^e.g. 55^. However, balance between Gα and Gβγ subunit expression may determine overall cAMP responses ^56^. As well, adenylyl cyclase isoforms can differ in their Gα sensitivities and expression levels can vary between immune cell types ^55^. Furthermore, G protein coupling “switches” have been reported for receptors in heterodimers ^57,58^. The particular profile and stoichiometries of interaction partners can give rise to differential signaling “units” between cell types ^reviewed in 59^. There are also various explanations for potential dual coupling within the same cell. Signalsome subcellular compartmentalization and/or sequestration of signaling components is one example ^reviewed in 60^; indeed subcellular distribution of CB_2_ can influence CB_2_ signaling patterns ^6^, and CB_2_ seems to have a particularly unique distribution in immune cells ^61–63^. Homologous signaling feedback upon CB_2_ activation which results in rapid desensitization of Gαs coupling with little or slower influence on Gα_i_ coupling such as via differential phosphorylation by G Protein-Coupled Receptor Kinases or PKA ^64–66^ might assist in explaining the observed cAMP signaling profile. Gβγ exhaustion/sequestration may also hinder Gα_s_ activation ^67,68^. It is also important to note, and intriguing regarding the downstream functional implications, that subcellular compartmentalization of G proteins and/or adenylyl cyclase subtypes could give rise to localized cAMP fluxes, implying that simultaneous effects of both cAMP increases and decreases are possible even when the net cAMP concentration is unchanged ^reviewed in 69^.

The ERK1/2 signaling cascade regulates cell cycle progression ^reviewed in 70^, immunological functions, including differentiation of T lymphocytes ^13^, polarization of CD4+ T lymphocytes ^71^, and regulation of cytokine network ^e.g. 72^, and is dysregulated in a number of disease states ^e.g. 23^. We find that human primary PBMC respond to CB_2_ activation by HU308 with rapid and transient stimulation of p-ERK. As this was completely sensitive to both Gβγ inhibitor gallein and Gα_i/o_/βγ inhibitor PTX, we conclude that ERK1/2 phosphorylation induced by HU308 is mediated by βγ dimers of Gα_i_βγ heterotrimeric proteins. Although in some systems gallein may inhibit GRK2 and possibly associated arrestin signaling ^73^, we do not know of any studies suggesting that PTX blocks arrestin-initiated signaling other than one report of a small potency shift for arrestin recruitment ^74^. Interestingly NF023, which we confirmed in cAMP assays inhibits Gα_i_, did not block the p-ERK response, indicating that this blocker specifically inhibits Gα subunits without influencing Gβγ activity as also observed previously ^75^. HU308 also induced phosphorylation of p38 and Akt. p-p38 was mediated by Gα_i/o_ subunits of G proteins (possibly via Src kinase) ^e.g. 76^, as it was completely blocked by Gα_i/o_/βγ inhibitor PTX but not by Gβγ inhibitor gallein, whereas p-Akt was mediated by Gα_i/o_–coupled βγ heterodimers (fully blocked by PTX and gallein).

In contrast with prior studies in human promyelocytic leukemia HL60 cells ^25^ and CB_2_-overexpressing HEK cells ^24^ we did not observe JNK phosphorylation. However, we note that activation of this pathway may be cell-context dependent given that in LPS-stimulated human monocytes JNK was inhibited by a CB_2_ ligand ^12^. Another explanation may be a different CB_2_-mediated functional profile (i.e. biased agonism) between HU308 and the previously studied ligands ^5,77^. There may also be signal-modifying influences of serum as has been shown for CB1 ^78^. The serum factors that circulating immune cells would be exposed to *in situ* are likely to influence signaling patterns ^79^. We therefore feel that inclusion of 10% fetal bovine serum is an important methodological improvement upon earlier studies carried out under low serum or serum-free conditions. However, we must acknowledge that this is of course not a perfect model of human serum from the matched donor. Some of the resistance to inclusion of serum in cannabinoid signaling studies is the potential for presence of endocannabinoids and precursor molecules which would complicate the interpretation of reductionist experiments ^e.g. 80^. However, careful attention to vehicle controls, the potency of HU308 effects, only subtle inverse agonist effects, and lack of effects of a neutral antagonist when applied alone, indicate that any influence in this regard was minimal to non-existent in our study.

cAMP/PKA, MAP kinases ERK and p38, and Akt/PKB all have the potential to induce phosphorylation of CREB ^reviewed in 49^. Immunological functions of p-CREB include anti-apoptotic effects on macrophages via p-38 ^81^, proliferative effects on T lymphocytes via p-38 ^82^ and p-ERK ^83^, and activation/proliferation of B lymphocytes via p-ERK ^84^. CB_2_-mediated p-CREB activation in primary PBMC is mediated primarily and to a similar degree by Gα_s_ subunits and non-Gα_i_-derived βγ dimers, and to a limited extent by Gα_i_ subunits. Though Gα_s_–coupled βγ is not widely appreciated to induce signaling events, at least a few examples exist in the literature ^e.g. 85,86^. p-CREB is unlikely to be downstream of pERK or p-Akt in our system given that these pathways were both associated with βγ from Gα_i_βγ heterotrimers, however Gα_i_-coupled p-p38 may be involved to a minor degree. One of the primary p-CREB activation pathways is likely via Gα_s_ leading to cAMP-activated PKA. This is interesting given that when measuring cAMP concentrations we did not detect any increase unless a Gα_i_ blocker was present. This implies that the consequences of Gα_i_ versus Gα_s_ activation on cAMP concentration are distinct, supporting theories that either CB_2_ is signaling differently between cell types or that subcellular compartmentalization can give rise to differential local cAMP concentrations with independent consequences for signaling.

Proceeding to study a physiologically relevant outcome, we detected a CB_2_-mediated induction of interleukins 6 and 10 at 6 and 12 h. Both were sensitive to NF449 > gallein > NF023, indicating that Gα_s_ is the main contributor to these responses. This pattern of influence of G protein inhibitors was similar to that for p-CREB, and given that p-CREB can induce transcription of IL-6 and IL-10 genes ^reviewed in 87^, it is very feasible that CREB phosphorylation is an important mediator of HU308-induced IL-6 and IL-10. Although in some instances IL-6 may be secreted from a pre-formed pool ^e.g. 88^ the observed induction of IL-6 by HU308 at 6 and 12 h likely involves its *de novo* synthesis ^e.g. 89^. To our knowledge, PBMC do not have intracellular granules of IL-10, and so it is most likely being synthesized *de novo* and then secreted. Kinetics of mRNA induction for IL-6 and IL-10 ^e.g. 90,91^ are consistent with a multi-hour lag in their production after CB_2_ activation.

Given that p-p38 was blocked by PTX and not gallein, this is a candidate for the NF023-sensitive (Gα_i_-requiring) component of IL-6/10 induction. p-p38 and p-Akt have also been linked to both IL-6 ^e.g. 92,93^ and IL-10 production ^e.g. 94,95^. Given the indicated involvement of CREB we were surprised to find that forskolin, a strong activator of signaling pathways downstream of adenylyl cyclases, did not influence cytokine secretion. We postulate that a co-stimulus enabled by CB_2_ activation but not by forskolin might be required for IL-6/10 induction ^reviewed in 49^. It is also plausible that the potential for subcellular compartmentalization and/or temporal integration of cAMP fluxes play critical roles. Application of specific signaling pathway inhibitors would be useful to more directly delineate the exact signaling pathways responsible.

The degree of cytokine induction we observed was small in comparison with the positive control utilized, LPS – a potent mediator of bacterial sepsis ^reviewed in 96^, which models systemic inflammation *in vitro*. Given that HU308 was ∼10 times less efficacious than LPS, it is unlikely that CB_2_ activation would result in dramatic systemic inflammatory reactions *in vivo*, but might rather elicit immunomodulatory effects with a potential for both pro- and anti-inflammatory functions – likely depending on the state of the immune system.

We did not observe modulation of cytokines IL-2, IL-4, IL-12, IL-13, IL-17A, MIP-1α, or TNF-α, likely because we avoided activation of PBMC with mitogenic/antigenic proliferative or cytokine-inductive stimuli to preserve their *in vivo* phenotype and hence could not detect any potential inhibitory effects of HU308.

We also noted that HU308 did not influence cell number of PBMC treated for 6 and 12 h, which is *prima facie* unexpected given prior reports of cannabinoid-induced cell death. However, these prior studies were carried out under completely different experimental paradigms ^17,e.g. 97,98^. Furthermore, this response has been found to be cell type- and ligand-dependent ^reviewed in 99^, and is not always mediated by cannabinoid receptors ^e.g. 100^.

IL-10 is an anti-inflammatory cytokine expressed by many immunocompetent cell types which regulates T-lymphocytic responses via direct mechanisms ^e.g. 101^ and by inhibiting effector functions of antigen-presenting cells ^e.g. 102,103^. IL-10 can also protect from excessive immunoreactivity by inhibiting overproduction of cytokines, *including* IL-6 which is widely described as pro-inflammatory. Typical IL-6 effects include induction of acute phase proteins, monocyte recruitment and macrophage infiltration, activation of plasma cells, and CD4+ lymphocyte differentiation ^reviewed in 104^. Based on these paradigms the combined release of IL-6 and IL-10 seems counter-intuitive. However, IL-6 can context-dependently exert anti-inflammatory effects, and indeed simultaneous production of IL-6 and IL-10 from mixed human primary immune cells has been observed previously ^105^.

The HU308-induced production of both IL-6 and IL-10 within the same time-frame, and with strikingly similar potencies and patterns of influence by inhibitors, seem to imply that CB_2_ either stimulates release of one cytokine which induces the second, or that the two cytokines are induced via the same signaling pathways. Considerable literature supports the first possibility; for example, IL-6 in concert with other cytokines can stimulate differentiation of naïve CD4+ T cells into IL-10-producing regulatory type 1 helper T (Tr1) or IL-17-producing T helper (T_H_-17) cells which have anti-inflammatory effects *in vivo* ^106,107^. Low dose recombinant IL-6 can also increase plasma IL-10 and have other anti-inflammatory effects in humans ^108^. However, given the acute time-frame within which we detected both IL-6 and IL-10 (by 6h), and slightly greater potency of the IL-10 response, it seems unlikely that in our system CB_2_ activation first induces IL-6 which subsequently leads to IL-10 secretion from a different cell type. It is possible that IL-6 and IL-10 were released independently from different cells, however, the near-identical profile of sensitivity to G protein inhibitors leads us to suspect that the most plausible scenario is that IL-6 and IL-10 are generated from the same cell type(s) in our system. Interestingly, these cytokines have coordinated mRNA regulation via the same nuclear site ^109^. IL-6 and IL-10 have classically been thought to be produced by T helper 2 (Th2) T lymphocytes ^110, reviewed in 111^, while concurrent production has been observed in monocytes, tumor-associated macrophages and dendritic cells ^112–114^. In future work it will be important to clarify the influence of CB_2_ activation on different immune subtypes and this knowledge will also assist in designing follow-on experiments to investigate other CB_2_-mediated immune cell functions.

There is some precedent for cannabinoid mediation of both IL-6 and IL-10. Endocannabinoids upregulated IL-6 in LPS-stimulated U937 cells ^80^ and in whole blood cultures ^115^. Cannabinoid-mediated IL-10 induction has been shown *in vivo* in mice ^116^, in murine LPS-stimulated macrophages ^11^, and in murine encephalitogenic T lymphocytes ^117^; the latter study also shows inhibition of IL-6 by cannabinoids, likely indicating species and/or cell type specificity. To our knowledge, CB_2_-mediated IL-10 production by human PBMC has never been shown before. Historically, there is a considerable body of data regarding the general effects of cannabis and THC on human immune function, though somewhat surprisingly, little has been reported on cytokine profiles ^reviewed in 118,119^. Increased serum IL-10 has been correlated with cannabis consumption ^120^, while elevated IL-6 and other pro-inflammatory markers were found in individuals with cannabis dependency ^121^. In a trial of THC in multiple sclerosis, cannabinoid treatments were found not to influence serum concentrations of IL-10 or CRP (a surrogate measure of IL-6), nor other measured cytokines, though sample sizes were small and variability between subjects large ^122^. Further, THC is a partial agonist at CB_2_ and so may have only subtle effects in comparison with full agonists. Only a few clinical trials of CB_2_-selective agonists have been undertaken and to our knowledge none have reported on cytokine profiles. We have also noted serious adverse effects from illicit synthetic cannabinoids, some of which are potent CB_2_ agonists ^123,124^. Although these adverse symptoms may be mediated by a non-cannabinoid-receptor target ^e.g. 125^, we wonder whether a potent and efficacious influence on cytokine balance and/or different cytokine profile arising from biased agonism via CB_2_ could contribute to deleterious effects.

The results of this study greatly expand our knowledge of CB_2_ signaling and functional implications in human primary PBMC under conditions closely preserving their *in situ* state. The discovery of unprecedented CB_2_ coupling to stimulatory (Gα_s_) proteins, their simultaneous activation alongside inhibitory (Gα_i_) proteins in a cAMP pathway, of Gα_s_-mediated effects on CREB phosphorylation, and of IL-6 and IL-10 induction, offer new perspectives on CB_2_ signaling and function which may be useful for drug discovery and investigations of CB_2_ signaling phenomena including functional selectivity. The effects of CB_2_ agonism we have observed in this study raise a number of questions and potential challenges, but reinforce the potential utility of CB_2_ ligands as immunomodulatory therapeutics.

## Methods

### Preparation of Human PBMC

Human primary PBMC were isolated from blood samples collected from 3 healthy volunteers (male and female, aged 22-26) after obtaining written informed consents in accordance with ethical approval from the University of Auckland Human Participants Ethics Committee (#014035).

Whole venous blood samples, 50mL per draw, 3-6 draws per donor (with each set of experiments performed on one draw from each donor) were collected into K_2_EDTA-coated BD Vacutainer® Blood Collection Tubes (Becton, Dickinson and Company), transferred to polypropylene tubes (Corning), and diluted 1.25 volumes of blood with 1 volume RPMI 1640 medium (HyClone) supplemented with 2mM L-Glutamine (Thermo Fisher Scientific).

PBMC were isolated by gradient centrifugation in Histopaque®-1077 (Sigma-Aldrich) in Greiner Leucosep® 50mL tubes (Greiner Bio-One) for 15 min at 800×g at room temperature. The collected PBMC were washed twice with RPMI by centrifugation for 10 min at 250×g, re-suspended in a culture medium – RPMI supplemented with 10% v/v fetal bovine serum (FBS; New Zealand-origin, Moregate Biotech) and 2mM L-Glutamine. Following isolation, PBMC were either used in assays immediately, or cryopreserved in 50% RPMI with 40% FBS and 10% DMSO and stored at -80°C for up to 12 months until later use. Cryopreservation did not affect CB_2_ expression (Fig. S9) or HU308 signaling responses of PBMC (Fig. S10).

### Cell Lines and Cell Culture

Human embryonic kidney (HEK) cell lines stably transfected with human CB_1_ (HEK-hCB1 ^126^) or human CB_2_ (HEK-hCB2 ^50^), or an untransfected wild type HEK Flp-in cell line (HEK wt, Invitrogen, R75007) were cultured in high-glucose DMEM (HyClone) with 10% v/v FBS, in the presence of Zeocin (InvivoGen) at 250µg/mL, Hygromycin B (Thermo Fisher Scientific) at 50µg/mL, or Zeocin at 100µg/mL, respectively.

### Whole Cell Radioligand Binding

Human primary PBMC or HEK cells were pelleted by centrifugation for 10 min at 250×*g* or 5 min at 125×*g* respectively, and re-suspended in binding medium (RPMI with 1mg/mL BSA, 2mM L-Glutamine, 25mM HEPES). PBMC (3×10^5^ cells per assay point), HEK-hCB_1_ or HEK-hCB_2_ (0.4×10^5^ diluted in 1.6×10^5^ HEK wt to avoid ligand depletion), or HEK wt (2×10^5^) cells were dispensed in Deep Well 96-well Plates (Gene Era Biotech) with reaction mix already present to produce a final 200µL reaction volume. The reaction mix included 5nM (final concentration) [^3^H]-CP55940 (PerkinElmer), and an appropriate displacer to measure non-specific displacement of the radioligand or a vehicle (DMSO) control to measure total binding of the radioligand. The displacer was either ACEA at 1µM (arachidonyl-2-chloroethylamide, synthesized from arachidonic acid as described in ^127^), or HU308 at 1µM (Tocris Bioscience).

Plates were incubated on the surface of a 15°C thermostatic bath for 3 h to reach steady-state, then placed on ice. Reactions were then rapidly filtered under vacuum through GF/C glass fiber filter-bottom 96-well microplates (Perkin Elmer, MA, USA) pre-treated with 0.1% polyethyleneimine (PEI, Sigma-Aldrich). After three washes with ice-cold wash buffer (50mM HEPES pH 7.4, 500mM NaCl, 1mg/mL BSA) filter plates were dried overnight, then sealed, and 50µL per well Irgasafe Plus scintillation fluid (Perkin Elmer) was dispensed. After 30 min, plates were counted in a Wallac MicroBeta® TriLux Liquid Scintillation Counter (Perkin Elmer). Each independent experiment was verified to have undergone <10% ligand depletion. CB_1_ and CB_2_ protein quantification was based on the maximum binding parameter B_max_, the concentration of binding sites for the radioligand ^128^, using the following formula based on equation 7.5.21 from ^129^:

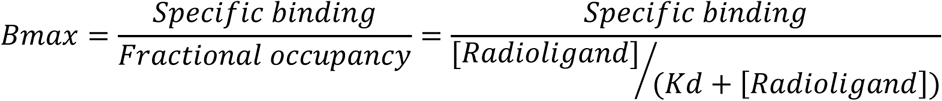

B_max_ were then converted to moles using the radioligand specific activity (as stated by the manufacturer), and then converted to units of fmol/mg of total protein added to an assay point (as measured by Bio-Rad DC Protein Assay Kit). Equilibrium dissociation constants (K_d_) for [^3^H]-CP55940 were measured by carrying out homologous competition binding under the same experimental conditions. pK_d_ values were: 7.94 ± 0.04 at CB_1_, 7.80 ± 0.05 at CB_2_ (*mean ± SEM, n=3*).

### cAMP Assay

PBMC were re-suspended in assay medium (RPMI with 2mM L-Glutamine, 25mM HEPES, 10% v/v FBS) and seeded at 1×10^5^ cells per well in 10µL into Falcon 384-well plates (Corning). All ligand and vehicle solutions were prepared at 2× concentrations in the assay medium, and dispensed 10µL per well in 384-well plates at requisite time points. After equilibration in a humidified atmosphere at 37°C and 5% CO_2_ for 1 h, cells were incubated with 10µM forskolin (canonical adenylyl cyclase activator, Tocris Bioscience) with or without stimulations of interest and/or vehicle (DMSO) controls. At the end of incubation, plates were rapidly cooled by placing on ice, and cells were lysed by adding 10µL per well of 3× concentrated ice-cold lysis buffer (formulation as per LANCE cAMP kit described below). The plates were placed on shakers at 500 RPM for 10 min at 4°C, and frozen at -80°C. The assays were performed without any phosphodiesterase inhibitors to allow for assessment of signaling kinetics.

cAMP detection was performed in 1/2-area white 96-well plates (PerkinElmer), using the LANCE cAMP Kit (PerkinElmer) with a high sensitivity method with 12µL per point lysate and total detection volume 24µL using 2× concentrated detection mix. Signal was detected on a CLARIOstar plate reader (BMG Labtech) using recommended TR-FRET settings. Assay sample cAMP concentrations were interpolated from standard curves. Data were normalized to vehicle (set as 0%) and forskolin (set as 100%) controls to allow compilation of data from independent experiments.

### ERK1/2 Phosphorylation Assay (p-ERK)

PBMC were pelleted by centrifugation for 10 min at 250×*g*, re-suspended in assay medium, and seeded at 1×10^5^ cells in 10µL per well into Falcon 384-well plates. Plates were incubated in a humidified atmosphere at 37°C and 5% CO_2_ for 55 min, and then transferred to the surface of a water bath at 37°C and incubated for 5 min prior to adding ligands or vehicle controls. Treatments were prepared at 2× concentrations in assay medium, and dispensed 10µL per well at requisite time points. At the end of incubation, plates were placed on ice, and cells were lysed by addition of 10µL per well of 3× concentrated ice-cold lysis buffer (formulation as per AlphaLISA kit described below). The plates were placed on shakers (500 RPM, 10 min at 4°C) and frozen at -80°C. p-ERK detection was performed using the AlphaLISA SureFire Ultra p-ERK 1/2 (Thr202/Tyr204) Assay Kit (PerkinElmer) per the manufacturer’s specifications in 1/2-area white 96-well plates. Signal was detected on a CLARIOstar plate reader with AlphaLISA-compatible filters. Counts were normalized to vehicle controls (100%) to allow compilation of data from independent experiments.

### Akt, p38, JNK, CREB Phosphorylation Assays

PBMC were treated and lysed under the same conditions as described for the p-ERK assay. Analytes were detected using AlphaLISA SureFire Ultra p-Akt 1/2/3 (Ser473), p-JNK 1/2/3 (Thr183/Tyr185), p-p38 MAPK (Thr180/Tyr182), and p-CREB (Ser133) assay kits as described above.

### Application of G Protein Inhibitors

In experiments with PTX (Sigma-Aldrich), cells were pelleted, re-suspended in the assay medium containing PTX 100 ng/mL or vehicle, and incubated for 5 h in a humidified atmosphere at 37°C and 5% CO_2_. After the incubation, the cells were counted and re-suspended in the same assay media with PTX or vehicle, equilibrated for 1 h as described for the signaling assays (bringing the total PTX incubation to 6 h), then ligands/vehicles added for signaling experiments. In gallein (Santa Cruz Biotechnology) or suramin derivatives NF023 (Abcam) and NF449 (Abcam) experiments, the inhibitors or their vehicles were added to cells at 10µM 30 min prior to adding ligands or vehicles. All ligand and vehicle solutions used in assays with inhibitors were prepared in assay media containing the inhibitors or vehicles, so that inhibitors or vehicle controls at 1× concentration were present for the entire duration of stimulations.

### Analysis of Cytokine Secretion

PBMC were seeded at 4.5×10^5^ cells per well in assay medium into 96 Well V-Bottom plates (Interlab), incubated for 30 min, if applicable pre-treated for 30 min with 1µM SR144528 (a kind gift from Roche Pharmaceuticals, Basel, Switzerland), 5µM WIN55212-3 (Tocris), 10µM G protein inhibitors (NF023, NF449, gallein), or vehicle control (DMSO), and subsequently treated with HU308 or vehicle, with or without forskolin and G protein inhibitors or antagonists if applicable, in 150µL final volume. After 12 h (multiplex screen) or 6 h (quantitative IL-6/IL-10 analysis) incubation in a humidified atmosphere at 37°C and 5% CO_2_, plates were centrifuged for 10 min at 250×*g*, and supernatants collected and frozen at -80°C. Remaining cell pellets were re-suspended and aliquots (in technical duplicate) diluted in Trypan blue for hemocytometer counting. Interleukins 2, 4, 6, 10, 12, 13, 17A, macrophage inflammatory protein (MIP-1α) and tumor necrosis factor (TNF-α) were detected using Cytometric Bead Array (CBA) technology (BD Biosciences) and assayed on an Accuri C6 flow cytometer (BD Biosciences) as described previously ^130^. For the multiplex screen, two concentrations of standard in the lower and middle range of standard curves (78 and 625 pg/mL) were utilized as internal controls and the mean fluorescent intensity (MFI) of each cytokine was calculated. Concentrations of IL-6 and IL-10 were interpolated using FCAP Array software (v. 3.1, BD Biosciences) from standard curves. Manufacturer stated limits of detection (LOD) were 11.2, 1.4, 1.6, 0.13, 7.9, 0.6, 0.3, 0.2, and 1.2 pg/mL for interleukins 2, 4, 6, 10, 12, 13, 17A, MIP-1α, and TNF-α, respectively.

### Data Analysis

All analysis was performed in GraphPad Prism Software (v. 8.0; GraphPad Software Inc.). Data is presented as mean ± standard error of the mean (SEM) from three independent experiments performed in triplicate, each with samples from a separate subject (three subjects in total). Concentration-response curves were obtained by fitting three-parameter nonlinear regression curves. Normalized data were utilized for statistical analysis of selected experiments (figures 2D-E, 4B, 5B, as described in figure legends). Otherwise, statistical analyses were performed on raw data. Repeated measures designs were utilized to reflect that each independent experiment was carried out on a separate subject. Normal distribution and equality of variance were verified with Shapiro-Wilk and Brown-Forsythe tests. If these tests did not pass, a decimal logarithm transformation was performed which enabled adherence to parametric test assumptions. If a statistically significant difference (*p < 0.05*) was detected in one- or two-way ANOVA, data were further analyzed using either Holm–Šídák (for comparisons to control) or Tukey (when comparisons between multiple conditions were of interest) *post hoc* tests with significance levels indicated graphically as: *p < 0.05* (*), *p < 0.01* (* *), *p < 0.001* (* * *), *non-significant* (ns).

## Supporting Information

Pre-incubation with forskolin has no effect on net cAMP kinetics in human primary PBMC; stimulatory cAMP signaling is detectable without forskolin; effects of HU308 on cAMP and p-ERK responses in PBMC are concentration-dependent; SR144528 is selective for CB_2_ over CB_1_; PBMC do not respond to a CB_1_-selective agonist ACEA in p-ERK assay; HU308 induces interleukins 6 and 10 but not IL-2, IL-4, IL-6, IL-10, IL-12, IL-13, IL-17A, MIP-1α or TNF-α in PBMC; HU308 has no effect on cell concentrations after 6 or 12 h of incubation; cryopreservation of PBMC has no effect on CB_2_ expression, nor p-ERK response.

## Acknowledgements

We thank John McGowan (University of Auckland, NZ) for synthesizing the ACEA. This research was supported by funding from the Auckland Medical Research Foundation and the Health Research Council NZ (to NG). YS was supported by Dean’s International Doctoral Scholarship, Faculty of Medical and Health Sciences, The University of Auckland. The Accuri C6 flow cytometer was purchased with funding from the NZ Lottery Health Board (to ESG).

