## Supplementary Figures for "Cannabinoid Receptor 2 (CB_2_) Signals via G-alpha-s and Induces IL-6 and IL-10 Cytokine Secretion in Human Primary Leukocytes"

**This PDF file includes:**

Figures S1 to S10

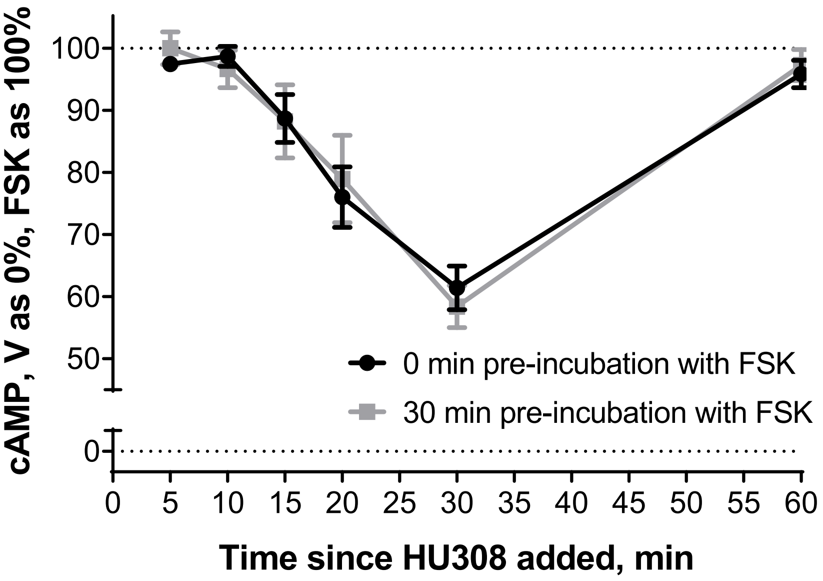


**Supplementary Fig. 1. cAMP signaling in human primary PBMC pre-incubated with forskolin.** Time-course of 1 μM HU308 with 10 μM forskolin (FSK) added at the same time (black) or after a 30-min pre-incubation with forskolin (grey). Data are normalized to vehicle control (0%) and forskolin alone (100%). Graph shows mean ± SEM for three independent experiments performed in technical triplicate, each on a separate subject (three subjects in total).

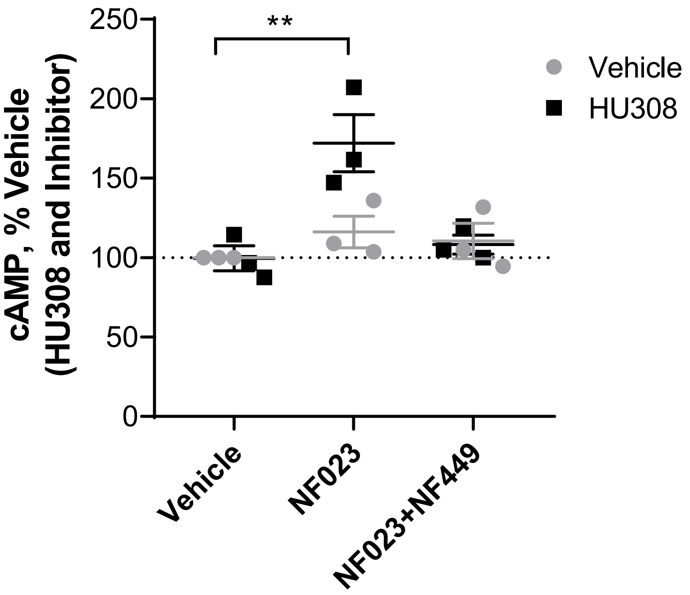


**Supplementary Fig. 2. Stimulatory cAMP signaling in human primary PBMC without forskolin.** PBMC were pre-treated for 30 min with a vehicle control, a $G\alpha_i$ inhibitor NF023 (10 μM), or a $G\alpha_s$ inhibitor NF449 (10 μM), and then HU308 (1 μM) was applied for 10 min (apparent signaling peak for stimulatory cAMP response). The responses are normalized to vehicle control (100%). All conditions were compared to the vehicle control. Only "NF023 + HU308" condition is significantly different from vehicle (*$P < 0.01$,* **), Tukey's multiple comparisons test. Graph shows independent experiment means (from technical triplicate) as well as overall mean ± SEM of these three independent experiments each performed with cells from a separate subject (three subjects in total). Note that NF023 reveals the stimulatory cAMP response, while NF449 blocks it.

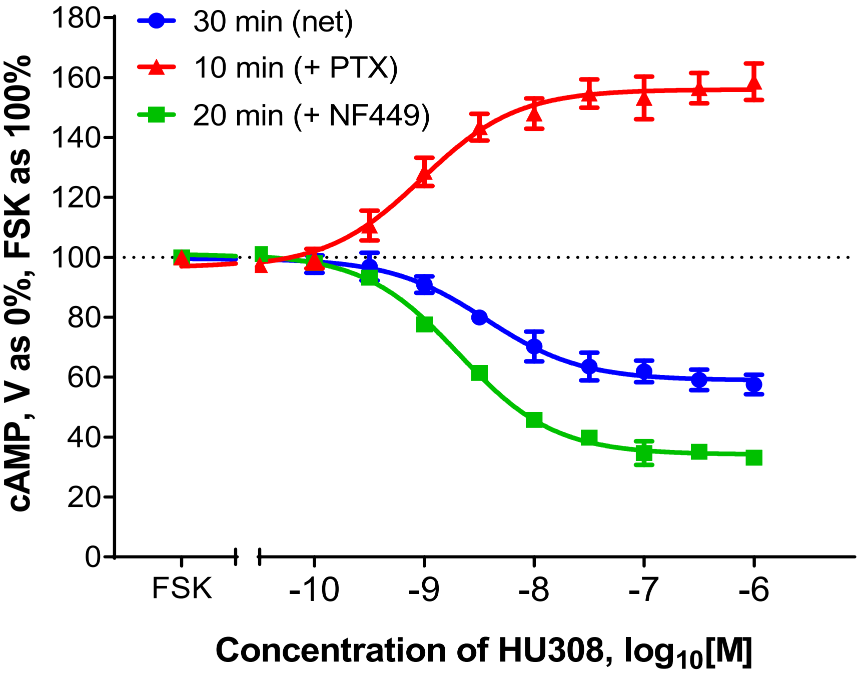


**Supplementary Fig. 3. Concentration-dependent effects of HU308 on cAMP responses in human primary PBMC.** Concentration-responses in the presence of 10 µM forskolin (FSK) and HU308 without inhibitors (●), or pre-treated with NF449 (■), or PTX (▲) and then stimulated with HU308 for 30, 10, or 20 min, respectively (at signaling peaks). Graph shows mean ± SEM for three experiments performed independently, each on a separate subject (three subjects in total).

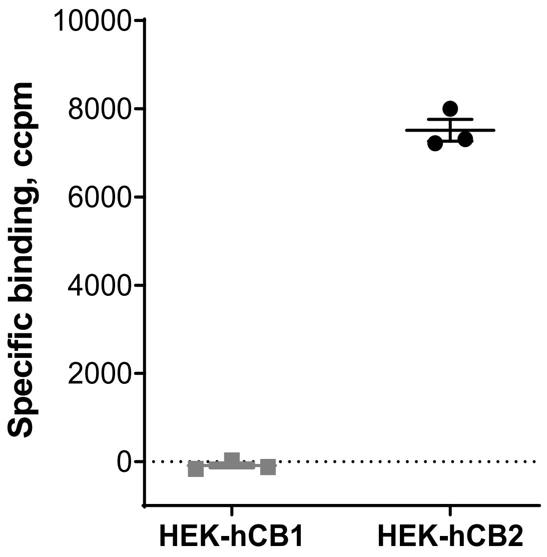


**Supplementary Fig. 4. Selectivity of SR144528 for $CB_2$ over $CB_1$.** HEK cells transfected with human $CB_1$ (HEK-h$CB_1$) or $CB_2$ (HEK-h$CB_2$) were incubated for 3 h at 15°C with [$^3$H]-CP55940 (5 nM) in the presence of a displacer SR144528 (1 μM, displaceable binding), or a vehicle control (total binding). The difference between total and displaceable binding was considered specific binding, plotted as ccpm – counts per minute corrected with a factory pre-set [$^3$H] detector normalization. HEK-h$CB_1$ is a positive control for $CB_1$, HEK-h$CB_2$ is a positive control for $CB_2$. Graph shows independent experiment means (from technical triplicate) as well as overall mean ± SEM of these three independent experiments each performed with cells from a separate subject (three subjects in total). Note absence of SR144528 displacement of [$^3$H]-CP55940 binding in $CB_1$-expressing cells.

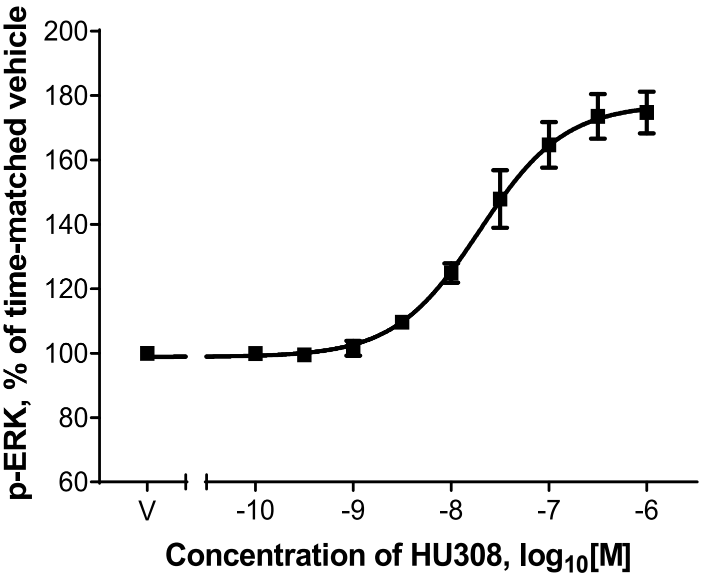


**Supplementary Fig. 5. Concentration-dependent effect of HU308 on p-ERK response in human primary PBMC.** HU308 concentration-response curve at 3 min, “V” indicates vehicle control. Graph shows mean ± SEM for three experiments performed independently, each with cells from a separate subject (three subjects in total).

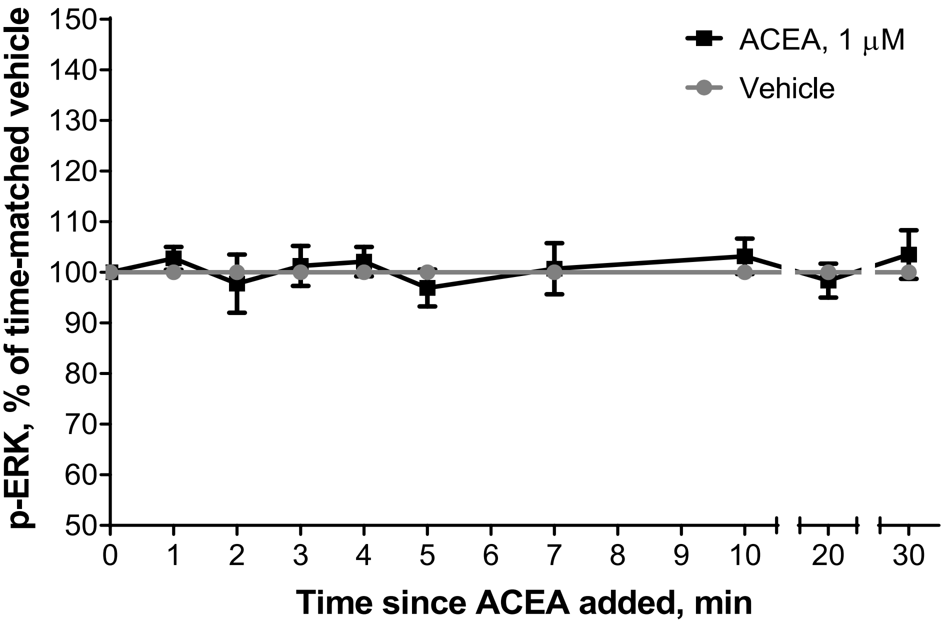


**Supplementary Fig. 6. p-ERK1/2 time-course in human primary PBMC.** Cells were treated with 1 μM ACEA (■) or a vehicle control (●). Responses are normalized to vehicle control at each time point. Graph shows mean ± SEM for three independent experiments performed in technical triplicate, each on a separate subject (three subjects in total). Note absence of ERK1/2 signaling in PBMC treated with a $CB_1$-selective agonist ACEA.

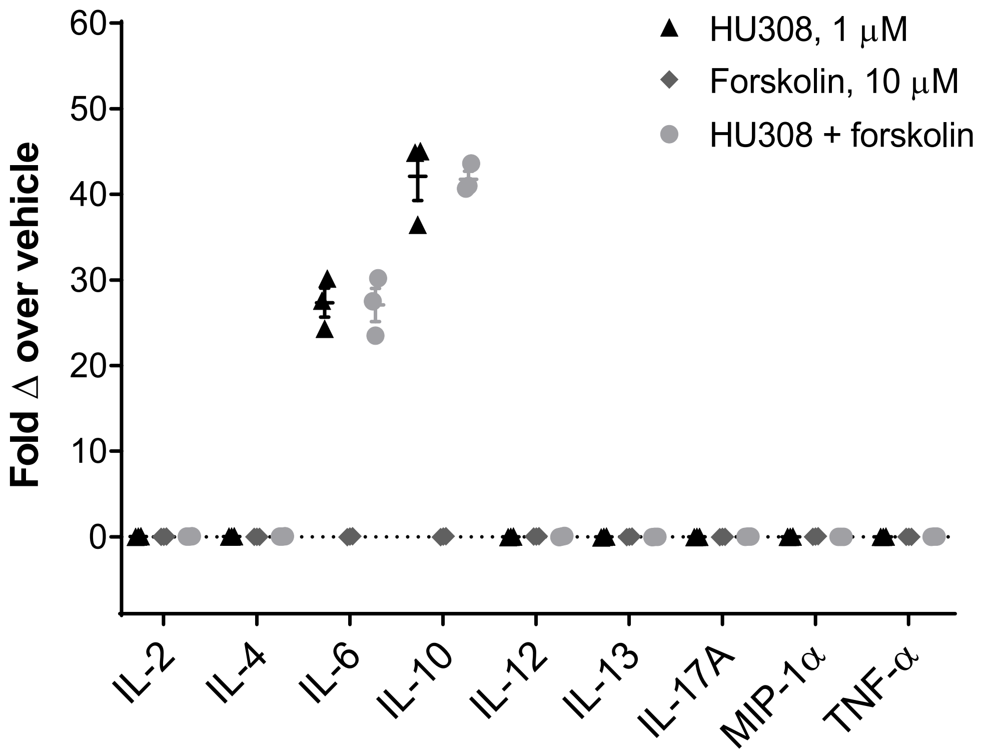


**Supplementary Fig 7. Induction of cytokine secretion by PBMC.** Cytokines in conditioned media from PBMC in response to HU308 (1 μM) in the presence or absence of forskolin (10 μM) after 12 h incubation. Graph shows independent experiment means (from technical triplicate) as well as overall mean ± SEM of these three independent experiments each performed with cells from a separate subject (three subjects in total).

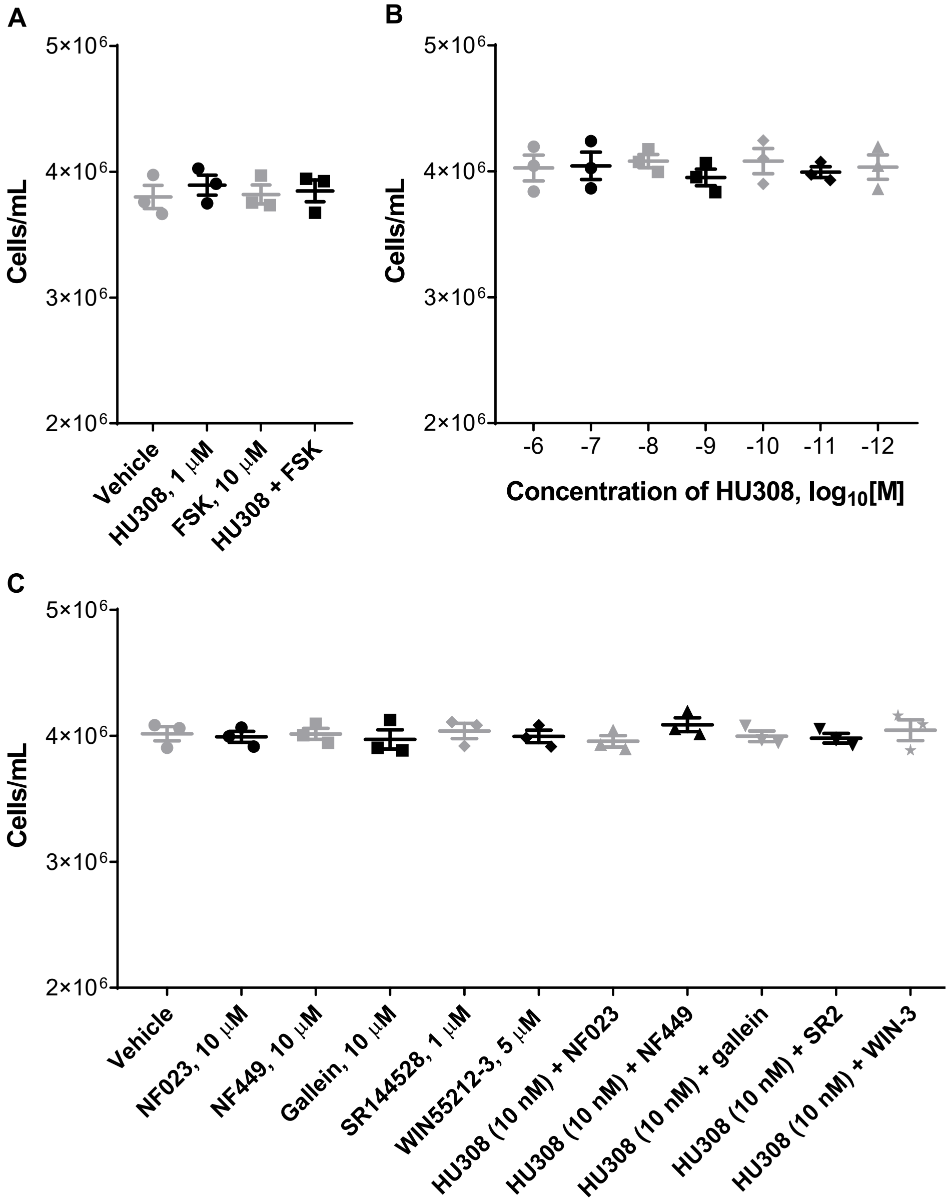


**Supplementary Fig. 8. Cell concentrations in PBMC samples used for cytokine secretion analysis.** PBMC were seeded at 4.5 × $10^5$ cells/well, and after incubations with the specified ligands or vehicle controls for 12 h (**A**) or 6 h (**B, C**) and supernatant collections, remaining cell pellets were re-suspended and counted in a hemocytometer. FSK – forskolin, SR2 – SR144528, WIN-3 – WIN55212-3. Note absence of cell concentration changes in all conditions. Graphs show independent experiment means (from technical triplicate) as well as overall mean ± SEM of these three independent experiments each performed with cells from a separate subject (three subjects in total).

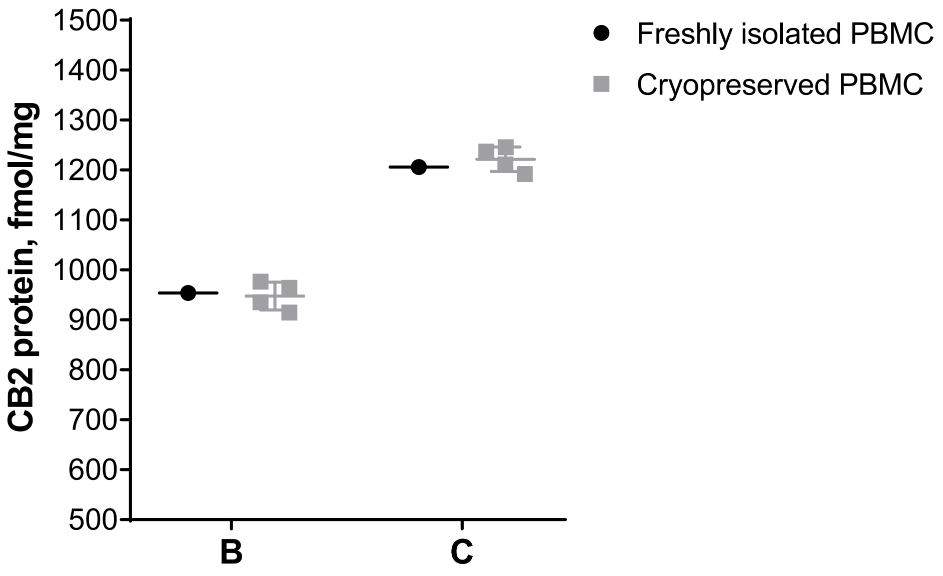


**Supplementary Fig. 9. $CB_2$ expression in freshly isolated and cryopreserved human primary PBMC.** Freshly isolated and cryopreserved (then resuscitated after 4 days, 2, 6, and 14 months) PBMC from subjects B and C were incubated for 3 h at 15°C with [$^3$H]-CP55940 (5 nM) in the presence of a displacer HU308 (1 μM, displaceable binding), or a vehicle control (total binding). The difference between total and HU308-displaceable binding was considered specific binding, and utilized to calculate total binding sites ($B_{max}$) and then converted to fmol/mg of total protein as described in methods. Graph shows means from 4 independent experiments performed in technical triplicate, with overall mean ± SD, with samples from two donors (B, C). Note that there is no difference in $CB_2$ expression in freshly isolated and cryopreserved for up to 14 months PBMC.

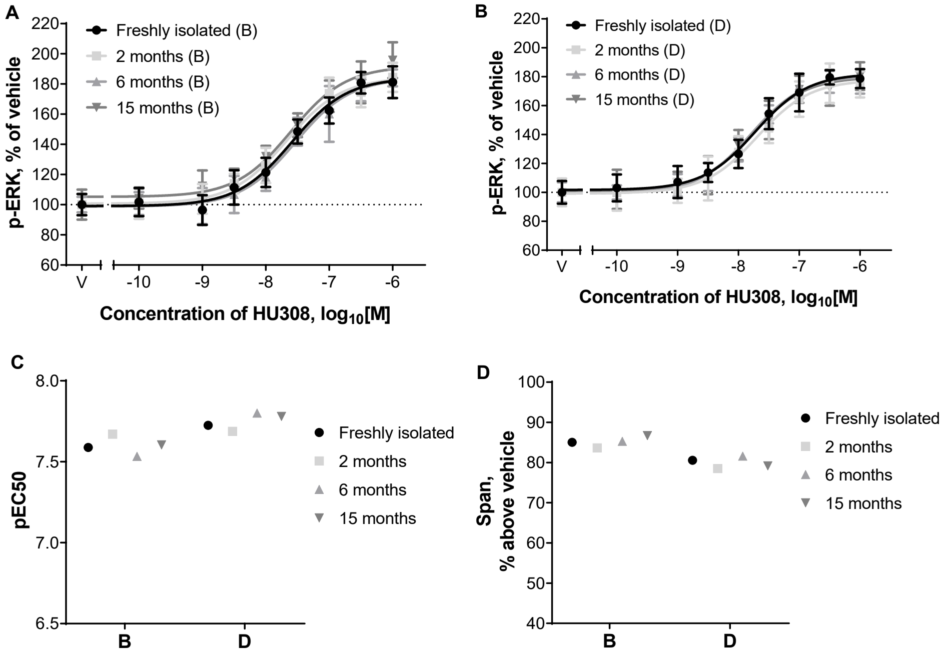


**Supplementary Fig. 10. p-ERK1/2 concentration-responses with freshly isolated and cryopreserved human primary PBMC. A** and **B**: freshly isolated and cryopreserved (then resuscitated after 2, 6, and 15 months) PBMC from subjects B and D, respectively, were treated with a concentration series of HU308 or a vehicle control (indicated as "V"); data is normalized to vehicle controls (100%); concentration-response curves were obtained by fitting three-parameter nonlinear regression curves (Hill slope constrained to 1) using GraphPad Prism Software. **C**. Potencies of HU308 in freshly isolated and cryopreserved PBMC from subjects B and D calculated from the curves A and B, respectively. **D**. Efficacies of HU308 in freshly isolated and cryopreserved PBMC from subjects B and D calculated as spans of the curves A and B, respectively, above vehicle controls. Graphs A and B show mean ± SD of four independent experiments performed in technical triplicate, each with PBMC samples from the same isolation, subjects B and D, respectively; graphs C and D show calculated parameters from each concentration-response curve (shown in A and B). Note that there is no difference in concentration-responses with freshly isolated and cryopreserved for up to 15 months PBMC.
